# Meta-analytic Review of Bacteria and Parasites in Elasmobranchs Provides Insight into Research Gaps

**DOI:** 10.1101/2024.09.27.614988

**Authors:** Alexis Modzelesky

## Abstract

Elasmobranchs have conservational and commercial importance. There is a need for a more complete understanding of their health, due to their ability to shape trophic webs and their money-making potential with fishers and ecotourists. Sharks have been known to influence the strength of food webs, including reef ecosystems where many fish species are harvested for food. The investigation of pathogenic agents and diseases in elasmobranchs has been biased to favor those natural enemies that inhabit the digestive system and epidermal surfaces. Certain groups of parasites such as cestodes, copepods, and monogeneans, were most often recorded in this review. The bacterial microbiota of elasmobranchs is currently being researched, but much information is lacking in the field, with the exception of certain well-known strains, such as members of the genus Vibrio. In addition, information on the 22 species of sharks included in this study is more in depth than that which can be found for the 11 ray species and three skates. A quantitative study was undertaken to investigate relationships between taxonomic Order and Family, as well as the relationship between diet and number of natural enemies with the use of the statistical software R. It was found that number of natural enemies present in an elasmobranch species correlates with the number of sources available. This first attempt at an exhaustive review of elasmobranch natural enemies from Florida waters includes over 400 pathogens as well as indicates the 14 most under-studied elasmobranch species.

## Introduction

Elasmobranchs have been known to influence the resiliency of food webs, including reef ecosystems where many fish species are harvested for food (NOAA 2001). Jameson et al. (1995) notes that some causes of population decline of reef organisms include anthropogenic stressors such as: overfishing, pollution, and increased sedimentation. Natural enemy information is critical to managing species of both conservation and commercial importance. Dobson and May (1987) attempted to predict how pathogens affect the dynamics of exploited fisheries and found that the ability for a pathogen to establish itself is dependent on the host’s current parasite density. Related to that same study, Winemiller (2005) created a model for predicting population resilience, production potential, and conformity to density dependent regulation in fish and found that his models can be useful in understanding traits as adaptations to environmental variation. He investigated three major life history strategies because this gives a more comprehensive view of adaptation to environmental variation than if one strategy was examined at a time. Burge et al. (2014) investigated another complicating factor for use within mathematical modeling of host-pathogen relationships: climate change. Burge et al. noted that marine ecosystems are among the most valuable and heavily used natural systems in the world, and provide many services, such as water filtration, food from fisheries and aquaculture, and tourism revenue. Effective management of species of interest requires understanding the factors that influence their populations, including their pathogens.

This study is a meta-analytic study of the parasites and diseases of elasmobranchs known to inhabit Florida coastal waters coupled with literature review. Although there are many reports of individual parasite taxa in elasmobranchs, at the time of writing no single study has compiled an exhaustive data search on natural enemy communities from as many countries as this study, nor has there been any attempt to explore such data using a meta-analytical approach. The scope of this research includes all parasites as well as viral, fungal, and bacterial infections found in selected elasmobranch species. The current study comprises information taken from elasmobranchs in both captive and wild situations. Pathogenic data comes from numerous countries, including: the United States, the United Kingdom, Mexico, Africa, Germany, Japan, Saudi Arabia, Russia, and Australia.

### Significance of Elasmobranchs

The importance of sharks as both apex and meso-predators has a large effect on ecosystem trophic webs. For example, blue shark Prionace glauca and white shark Carcharodon carcharias are mentioned as species known to have large distributions and can have profound effects on trophic webs and shaping them, especially in the open ocean (Wood et al. 2010). Wood et al. (2010) mentions that the removal of fish from the world’s oceans can be driving a long term and global decline in their parasites as well. In this same study the authors go on to say, “Because fishing reduces the density of fish (reducing transmission efficiency of directly transmitted parasites), this selectively removes large fish (which tend to carry more parasites than small fish), and reduces food web complexity (reducing transmission efficiency of trophically transmitted parasites), therefore the removal of fish from the world’s oceans over the course of hundreds of years may be driving a long term, global decline in fish parasites.” From a trophic perspective, Heithaus et al. (2008) investigated ecological consequences when marine top predator populations decline. While the reported effects varied, the loss of apex predators such as large sharks were often found to have large cascading effects. For example, with an increase in the number of cownose rays, Rhinoptera bonasus, the number of shark catches decreases. This increase in the population of cownose rays can cause a collapse in certain fisheries such as those for bivalves (an important fisheries resource), but this influence has not been studied in depth.

### Phylogeny

Members of Chondrichthyes - cartilaginous fishes - include sharks, rays, skates, and chimaeras. Camhi et al. (2008) noted sharks and rays are some of the oldest vertebrates in the oceans, with elasmobranchs surviving for more than 400 million years. About 360 million years ago, ancestral sharks diversified extensively (Bright 2011). Vélez-Zuazo and Agnarsson (2011) studied the phylogeny of shark and ray species in an attempt to resolve phylogenetic discrepancies due the creation of tree models based on morphology rather than molecular evidence. Vélez-Zuazo and Agnarsson (2011) note that as the field of molecular biology advances, the previous tree models that were based on morphology will be updated based on molecular and genetic information. There are some agreements on major classifications, such as taxonomic Order. In the present meta-analytical study, the Orders included are: Carcharhiniformes, Orectolobiformes, Lamniformes, Myliobatiformes, Rajiformes, Rhinopristiformes, and Squatiniformes. Vélez-Zuazo and Agnarsson go on to say that Carcharhiniformes were considered a sister taxa to Lamniformes, supported by morphology and molecular evidence, as well as sister to Orectolobiformes (Naylor et al. 2005; Compagno, 1973; de Carvalho 1996; Douady et al., 2003; Human et al., 2006; Heinicke et al., 2009; Mallatt and Winchell 2007; as cited by Vélez-Zuazo and Agnarsson 2011).

Taxonomic Family is considered more controversial. Within the Carcharhiniformes, for example, the families Scyliorhinidae and Triakidae appear to be paraphyletic (Iglésias et al., 2005; Human et al., 2006; Lopez et al., 2006 as cited by Vélez-Zuazo) and Odontaspididae and Carcharhinidae, often considered a single family (reviewed in Shimada et al., 2009), formed divergent branches in Lamniformes (Martin et al., 2002). Within the Squaliformes, the monophyly and relationships among most of the families were more or less completely unresolved until somewhat recently (Klug and Kriwet, 2010). While the current study does not attempt to parse phylogeny, the families included in this study are: Carcharhinidae, Cetorhinidae, Dasyatidae, Ginglymostomatidae, Gymnuridae, Lamnidae, Mobulidae, Myliobatidae, Narcinidae, Pristidae, Rajidae, Rhincodontidae, Rhinobatidae, Scyliorhinidae, Sphyrnidae, Squatinidae, Torpedinidae, Triakidae.

### Species Included in this Study

- Carcharhinus acronotus (Blacknose shark)
- Carcharhinus brevipinna (Spinner Shark)
- Carcharhinus isodon (Finetooth Shark)
- Carcharhinus leucas (Bull Shark)
- Carcharhinus limbatus (Blacktip Shark)
- Carcharhinus plumbeus (Sandbar Shark)
- Carcharhinus signatus (Night Shark)
- Carcharodon carcharias (White Shark)
- Cetorhinus maximus (Basking Shark)
- Galeocerdo cuvier (Tiger Shark)
- Ginglymostoma cirratum (Nurse Shark)
- Isurus oxyrinchus (Shortfin Mako)
- Mustelus canis (Smooth dogfish/ dusky smooth hound)
- Negaprion brevirostris (Lemon Shark)
- Prionace glauca (Blue Shark)
- Rhincodon typus (Whale Shark)
- Rhizoprionodon terraenovae (Atlantic Sharpnose Shark)
- Scyliorhinus retifer (Chain catshark/chain dogfish)
- Sphyrna lewini (Scalloped Hammerhead)
- Sphyrna mokarran (Great Hammerhead)
- Sphyrna tiburo (Bonnethead)
- Squatina dumeril (Atlantic Angelshark/ Sand Devil/ Devil Shark)
- Aetobatus narinari (Spotted Eagle Ray)
- Dasyatis americana (Southern Stingray)
- Dasyatis sabina (Atlantic Stingray)
- Dasyatis say (Bluntnose stingray)
- Gymnura micrura (Smooth butterfly ray/lesser butterfly ray)
- Manta birostris (Manta)
- Narcine bancroftii (Lesser electric ray)
- Pristis pectinata (Smalltooth sawfish/wide sawfish)
- Rhinobatos lentiginosus (Atlantic Guitarfish/Freckled Guitarfish)
- Rhinoptera bonasus (Cownose ray)
- Tetronarce nobiliana (Atlantic torpedo)
- Dipturus laevis (Barndoor skate)
- Raja eglanteria (Clearnose skate)
- Raja texana (Roundel skate/Texas Clearnose Skate)

### Pathogen Background

Natural elasmobranch populations are exposed to a wide range of viral, bacterial, and parasitic microbiota due to their diverse distributions. While some natural enemies have limited pathologies, others can be so pathogenic as to regulate their hosts’ populations, such as the bacterial genus Vibrio found in teleost fishes (Austin and Austin 2007). Parasites, bacteria, and viruses are ubiquitous, however, strains of pathogenic microorganisms can be very damaging when they enter the body of an elasmobranch. Both acute diseases of individuals and chronic outbreaks are possible, especially in fish farms (Austin and Austin 2016) and other captive situations. In many cases, a lesion or soft tissue damage is the perfect entryway for bacteria to infiltrate the host and begin causing immune responses.

### Current Work on Natural Enemies

Presently, there are few encompassing studies of the natural enemy communities of elasmobranchs. Garner (2013) published a summary of elasmobranch diseases, including several caused by parasites and other infectious agents, with special emphasis on microbial pathogens and their associated histopathology. Terrell (2004) and Goertz (2004) compiled a general list of pathogens for elasmobranchs commonly held in captivity, but the emphasis was on methods to treat some of the more prevalent infections (their reviews were part of a larger husbandry guide that listed more than 300 “taxa” of potential infectious agents). Cheung (1993) compiled a list of hundreds of elasmobranch species globally and their reported parasites, including some species covered in the present study; however, that study included some misnamed elasmobranchs and inconsistently-identified parasites.

An example of a commonly studied elasmobranch ectoparasite is the copepod (Benz and Deets 1986, Benz and Deets 1988, Benz 1989, Benz and Dupre 1987, Benz et al 2001, Dippenaar et al. 2008, and many others). In contrast, the information on bacterial inhabitants of elasmobranchs can be lacking, with the exception of the genus Vibrio being well documented (Austin and Austin 2007).

Some bodies of water are more studied than others, for example, areas found to be common fisheries would have the most available data. There is a plethora of information for elasmobranch parasites from areas such as the Sea of Japan and areas near Southeast Asia (Cheung 1993). Luckily, South Florida is a popular fishing destination and the universities have ample access to perform research on many of the resident elasmobranch species, with elasmobranch-related papers used in this study focusing on the Florida Keys and Gulf of Mexico areas (Caira and Gavarrino 1990, Cheung 1993, Karns et al. 2017).

The host / natural enemy list compiled in the present study will be of interest to resource managers and wildlife biologists who study elasmobranchs; providing much-needed information on potential causes of morbidity and mortality in local shark populations. Knowledge gaps focusing on individual elasmobranch species will be investigated and identified, which may open the door to further research by researchers.

## Materials and Methods

### Species selection

A search of online resources identified 36 elasmobranchs that frequent Florida waters. This included online repositories and websites of government organizations such as the Florida Fish and Wildlife Commission (FWC), the National Oceanic and Atmospheric Administration (NOAA), the International Union for Conservation of Nature (IUCN), as well as museum websites, and many peer reviewed articles posted to Google Scholar. Published accounts of the viruses, bacteria, and parasites of these species were obtained from online databases (Web of Science and EBSCO Host), and the host-parasite database compiled by the Museum of Natural History in London was used to cross-reference any sources already found, and provided an additional 30+ sources of information used in the study’s dataset. The following keywords were used in the database search: parasit*, bacter*, diseas*, patho*, elasmobranch*, as well as all the scientific names of the 36 shark, ray, and skate species. The IUCN, FishBase, and WoRMS (World Register of Marine Species) websites were also used to resolve any ambiguity of common names in elasmobranchs with large or global distributions. Additional databases such as Zoological Record were consulted, as well as any ‘grey’ literature sources available online, including unpublished government surveys and academic theses and dissertations. It is worth noting that some ambiguity in identifying specimens, whether elasmobranch or pathogen, may be due to: lack of quality specimens for examination as was the case for Beveridge (1993) with brittle parasites and low number of specimens in museum or personal collections or inability to identify the specimen caught by another party such as for Cheung (1993). If the species cannot be named due to degradation or for any reason, this is very important as one of the goals of the present meta-analytical study was to attempt an exhaustive search of natural enemies. In addition, the name may be considered *nomina dubia/species inquirenda* or the taxonomic category may be under review (Beveridge and Campbell 1988, Beveridge and Campbell 1998, Beveridge and Campbell 2010).

### Data analysis

A database was assembled in a spreadsheet as a matrix of elasmobranch species names with a complete presence/absence inventory of their natural enemies. Wherever possible, the abundance or other measure of infection intensity or prevalence was recorded and summarized; however, the current study is primarily concerned with which pathogens are present, so most analyses used presence-absence approaches.

These data were analyzed using two different strategies: first, the host-natural enemy data was used in multivariate analyses to explore how host factors shaped overall community structure; second, these data were then used to calculate more general indices of overall natural enemy diversity (i.e., the number of natural enemies recorded from each elasmobranch species in the literature), which were then tested against host factors (see below) using univariate statistical approaches such as regression.

Elasmobranch life history parameters were also determined from the literature and Fishbase.org, to establish a pool of possible predictor variables shaping natural enemy communities. Factors include size, preferred temperatures by the selected species with behavior and biological range information, diet, biometrics such as age at maturity, and average lifespan. Data on the feeding ecology of each elasmobranch species was collected from published accounts summarized in FishBase (Froese and Pauly 2017). Data was also collected on trophic level. Presence-absence data was imported into R studio (version 1.1.447, vegan package add-on).

### Univariate Approaches

Regression was used to assess the effects of univariate host factors (number of prey items, number of reports) on overall number of natural enemies.

### Multivariate Approaches

First, Sorensen similarity indices were calculated for their natural enemy communities in each elasmobranch species. The Sorensen index is a metric of community similarity that is specifically structured for use with presence-absence data. Sorensen indices were used to establish a triangular matrix of natural enemy community similarities among elasmobranch species. This triangular matrix was then used to statistically compare pathogen community structure in different elasmobranch taxa using an NMDS (Non-metric multidimensional scale) to search for possible patterns based on the presence or absence data. Clustering by taxonomic Family and Order were used as well in R, using hierarchical clusters with the hclust command. Hierarchical clustering in R is done using complete linkage; this method defines the cluster distance between two clusters to be the maximum distance between their individual components.

## Results

### Database Results

Much of the information on elasmobranch natural enemies has been produced by a select number of authors and some of the natural enemy types have been more widely studied (e.g., cestodes, with hundreds of species described in the literature). Table 1 (below) lists the 36 elasmobranch species by Family, their scientific name, the geographic region from which the species was recorded, and the associated sources. Some sources recorded a species as having been cataloged in more than one geographic region, and sources which included both wild and captive elasmobranchs. In the situation that information used in a source was from a museum collection that did not identify geographic region, Other (O) was indicated.

**Table 1.**
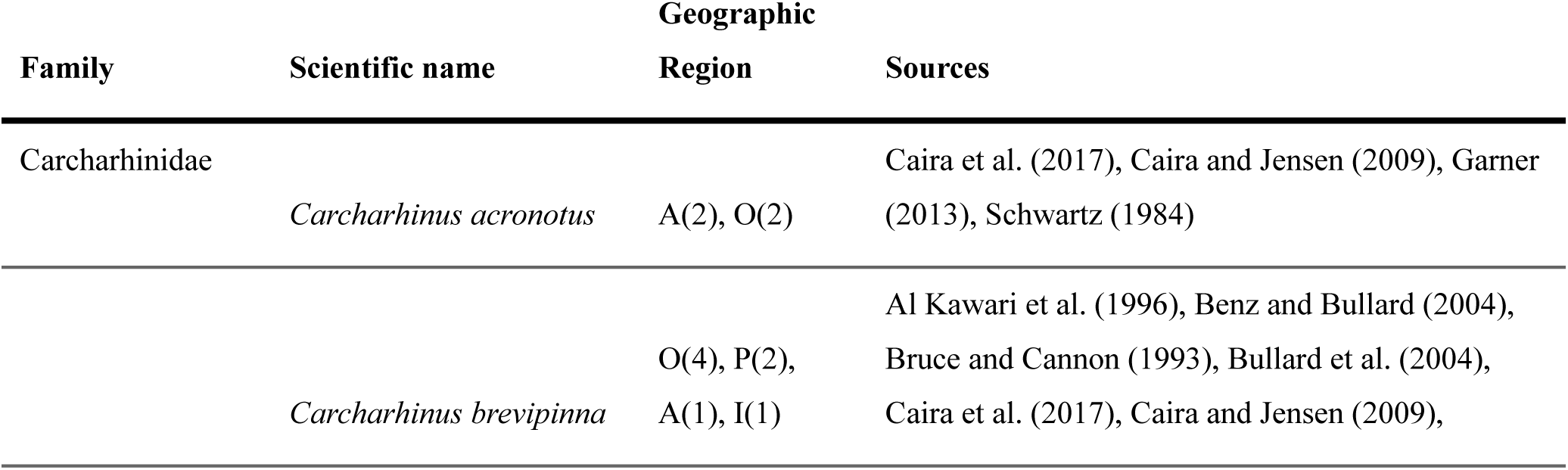
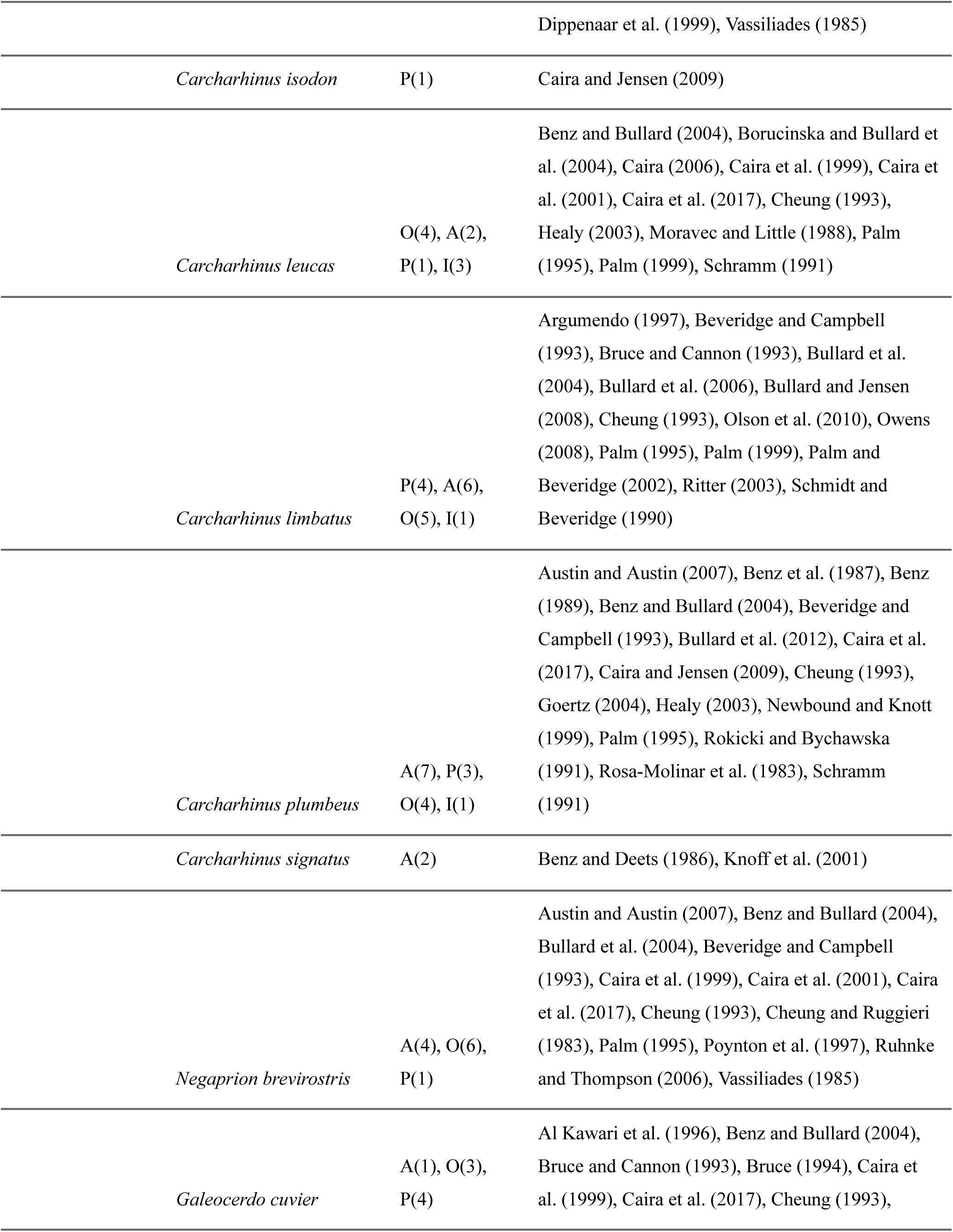
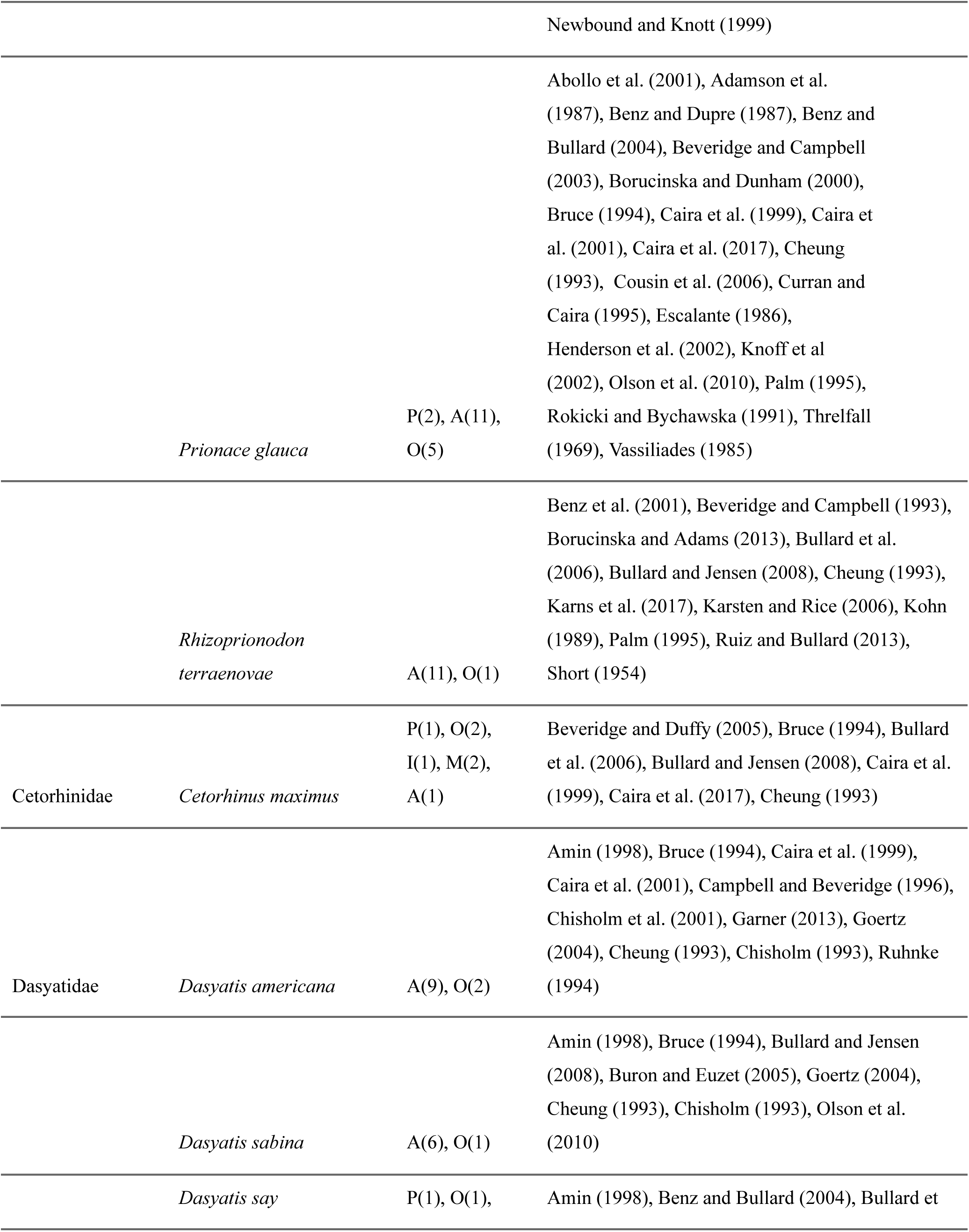
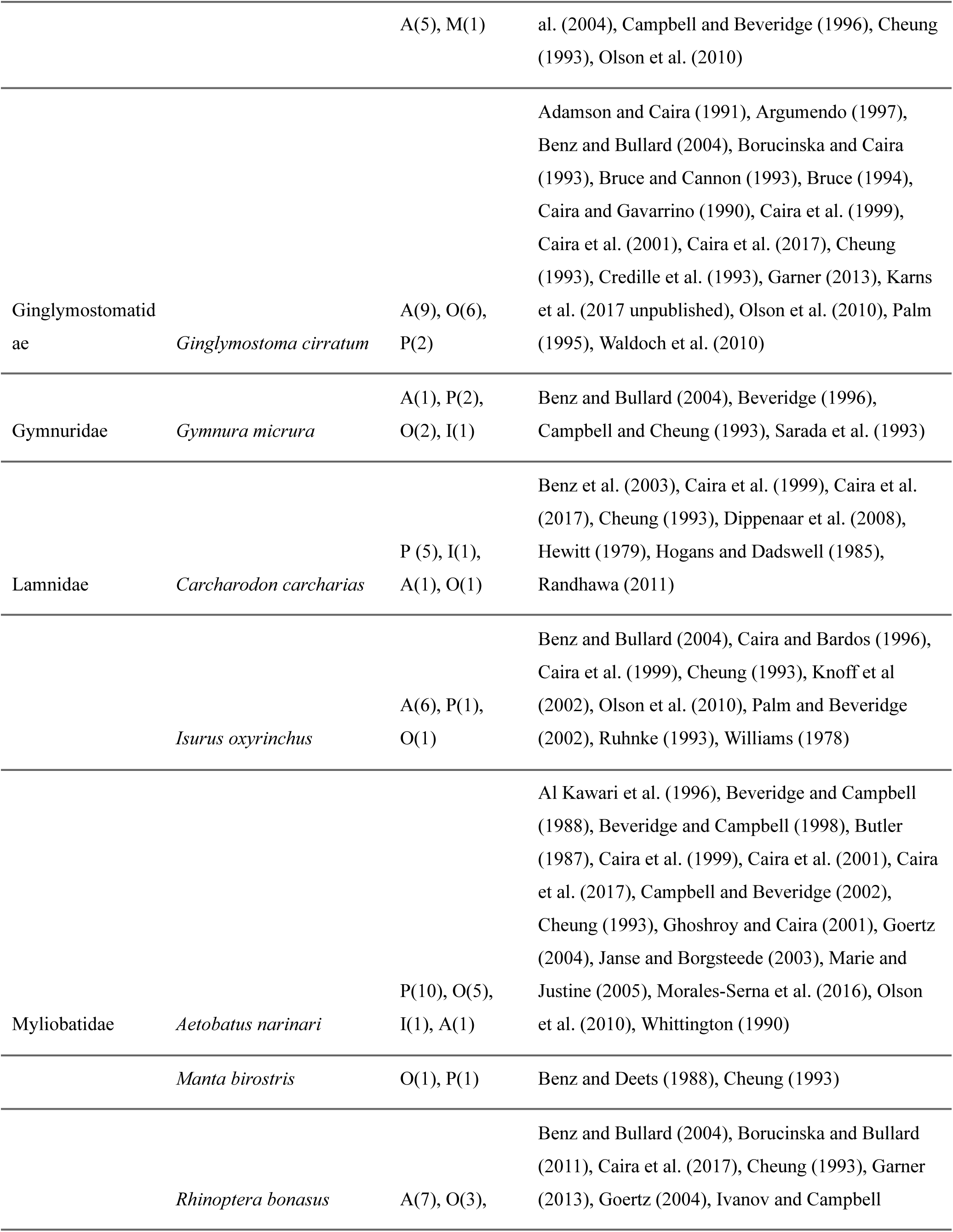
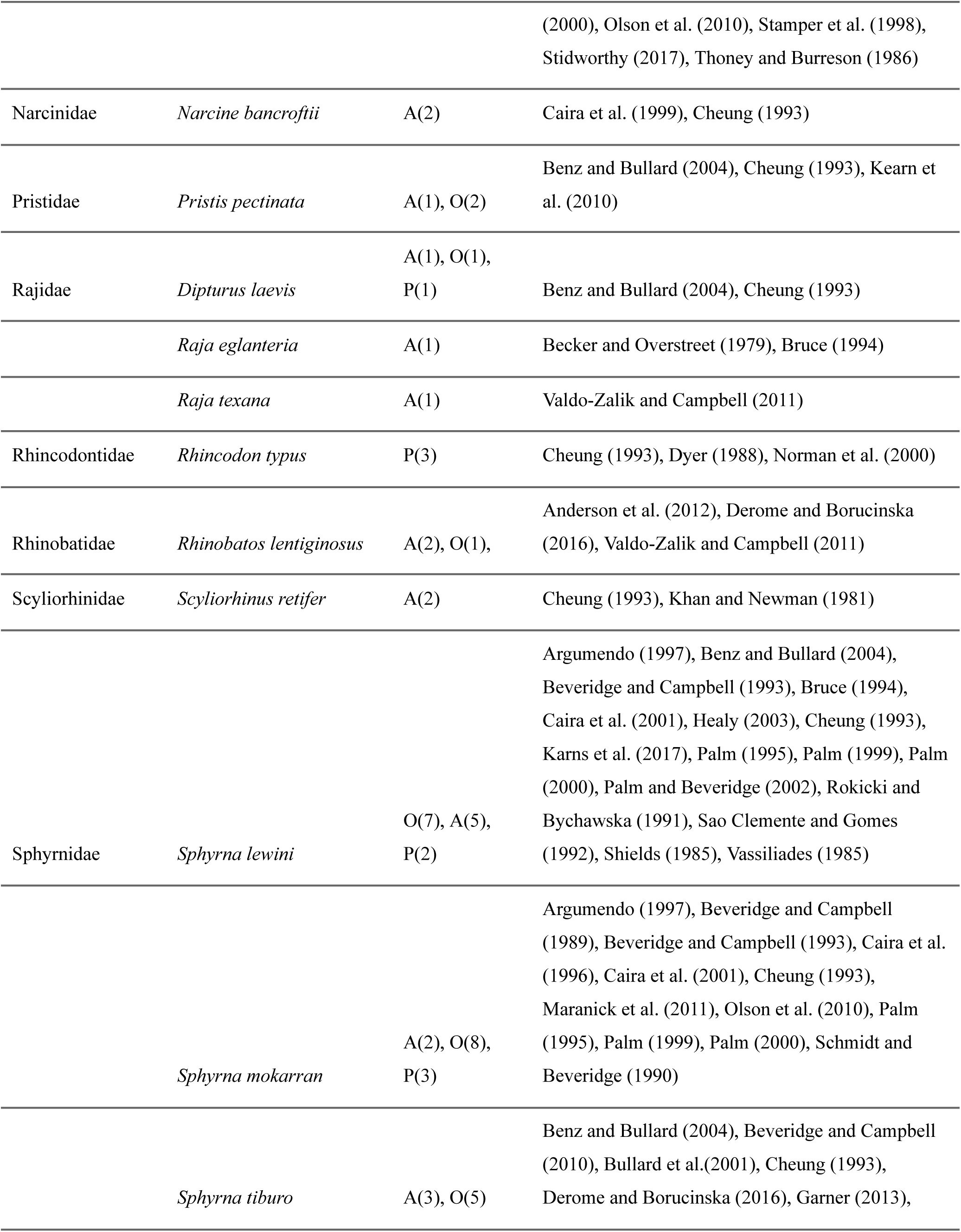
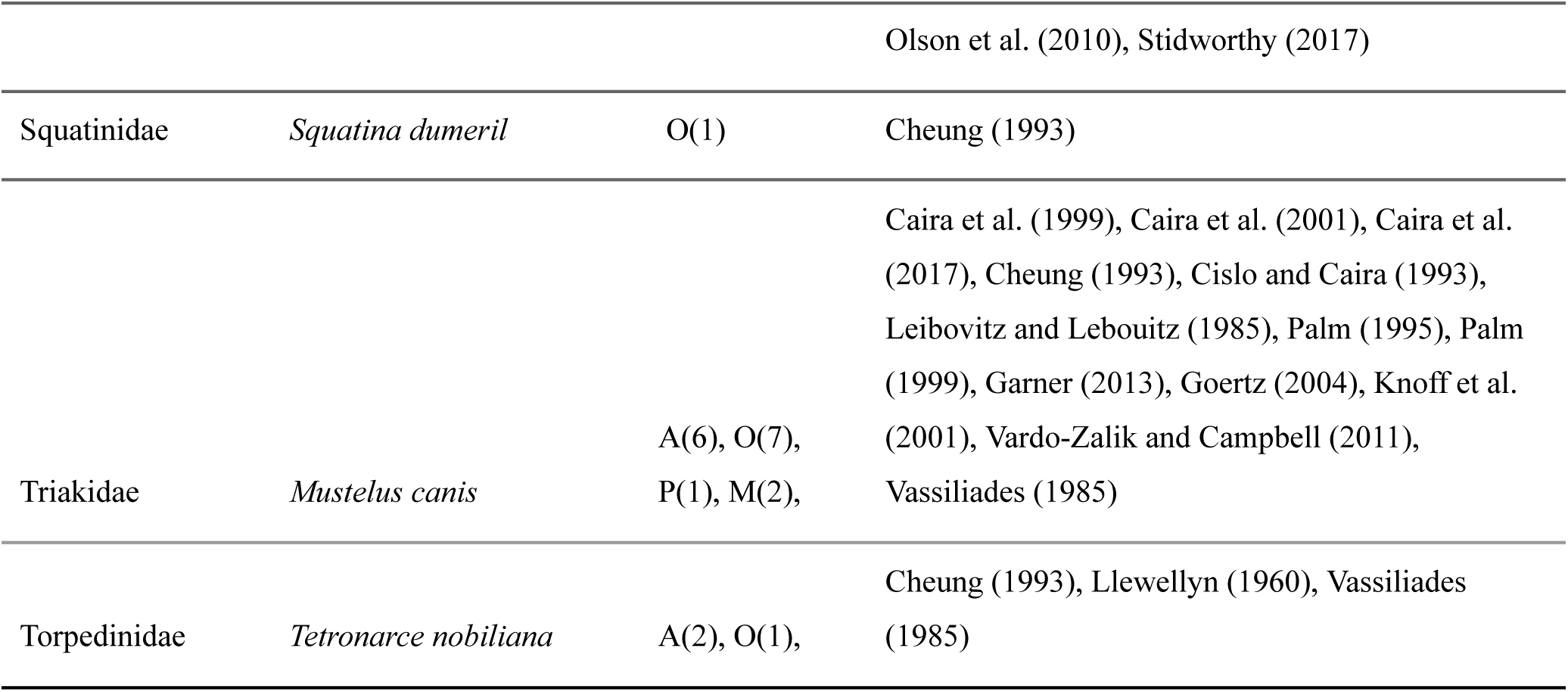
Peer-reviewed and other scientific literature found for 36 target species of elasmobranchs in a preliminary search. For geographic regions, A=Atlantic Ocean, P=Pacific Ocean, I=Indian Ocean, M=Mediterranean Sea, and O=other; numbers in parentheses indicate the number of individual references.

| Family | Scientific name | Geographic<br>Region | Sources |
| --- | --- | --- | --- |
| Carcharhinidae | *Carcharhinus acronotus* | A(2), O(2) | Caira et al. (2017), Caira and Jensen (2009), Garner (2013), Schwartz (1984) |
|  | *Carcharhinus brevipinna* | O(4), P(2),<br>A(1), I(1) | Al Kawari et al. (1996), Benz and Bullard (2004),<br>Bruce and Cannon (1993), Bullard et al. (2004),<br>Caira et al. (2017), Caira and Jensen (2009), |

|  |  |  |
| --- | --- | --- |
|  |  | Dippenaar et al. (1999), Vassiliades (1985) |
| *Carcharhinus isodon* | P(1) | Caira and Jensen (2009) |
| *Carcharhinus leucas* | O(4), A(2),<br>P(1), I(3) | Benz and Bullard (2004), Borucinska and Bullard et al. (2004), Caira (2006), Caira et al. (1999), Caira et al. (2001), Caira et al. (2017), Cheung (1993), Healy (2003), Moravec and Little (1988), Palm (1995), Palm (1999), Schramm (1991) |
| *Carcharhinus limbatus* | P(4), A(6),<br>O(5), I(1) | Argumendo (1997), Beveridge and Campbell (1993), Bruce and Cannon (1993), Bullard et al. (2004), Bullard et al. (2006), Bullard and Jensen (2008), Cheung (1993), Olson et al. (2010), Owens (2008), Palm (1995), Palm (1999), Palm and Beveridge (2002), Ritter (2003), Schmidt and Beveridge (1990) |
| *Carcharhinus plumbeus* | A(7), P(3),<br>O(4), I(1) | Austin and Austin (2007), Benz et al. (1987), Benz (1989), Benz and Bullard (2004), Beveridge and Campbell (1993), Bullard et al. (2012), Caira et al. (2017), Caira and Jensen (2009), Cheung (1993), Goertz (2004), Healy (2003), Newbound and Knott (1999), Palm (1995), Rokicki and Bychawska (1991), Rosa-Molinar et al. (1983), Schramm (1991) |
| *Carcharhinus signatus* | A(2) | Benz and Deets (1986), Knoff et al. (2001) |
| *Negaprion brevirostris* | A(4), O(6),<br>P(1) | Austin and Austin (2007), Benz and Bullard (2004), Bullard et al. (2004), Beveridge and Campbell (1993), Caira et al. (1999), Caira et al. (2001), Caira et al. (2017), Cheung (1993), Cheung and Ruggieri (1983), Palm (1995), Poynton et al. (1997), Ruhnke and Thompson (2006), Vassiliades (1985) |
| *Galeocerdo cuvier* | A(1), O(3),<br>P(4) | Al Kawari et al. (1996), Benz and Bullard (2004), Bruce and Cannon (1993), Bruce (1994), Caira et al. (1999), Caira et al. (2017), Cheung (1993), |
|  |  |  | Newbound and Knott (1999) |
|  | *Prionace glauca* | P(2), A(11),<br>O(5) | Abollo et al. (2001), Adamson et al. (1987), Benz and Dupre (1987), Benz and Bullard (2004), Beveridge and Campbell (2003), Borucinska and Dunham (2000), Bruce (1994), Caira et al. (1999), Caira et al. (2001), Caira et al. (2017), Cheung (1993), Cousin et al. (2006), Curran and Caira (1995), Escalante (1986), Henderson et al. (2002), Knoff et al (2002), Olson et al. (2010), Palm (1995), Rokicki and Bychawska (1991), Threlfall (1969), Vassiliades (1985) |
|  | *Rhizoprionodon terraenovae* | A(11), O(1) | Benz et al. (2001), Beveridge and Campbell (1993), Borucinska and Adams (2013), Bullard et al. (2006), Bullard and Jensen (2008), Cheung (1993), Karns et al. (2017), Karsten and Rice (2006), Kohn (1989), Palm (1995), Ruiz and Bullard (2013), Short (1954) |
| Cetorhinidae | *Cetorhinus maximus* | P(1), O(2),<br>I(1), M(2),<br>A(1) | Beveridge and Duffy (2005), Bruce (1994), Bullard et al. (2006), Bullard and Jensen (2008), Caira et al. (1999), Caira et al. (2017), Cheung (1993) |
| Dasyatidae | *Dasyatis americana* | A(9), O(2) | Amin (1998), Bruce (1994), Caira et al. (1999), Caira et al. (2001), Campbell and Beveridge (1996), Chisholm et al. (2001), Garner (2013), Goertz (2004), Cheung (1993), Chisholm (1993), Ruhnke (1994) |
|  | *Dasyatis sabina* | A(6), O(1) | Amin (1998), Bruce (1994), Bullard and Jensen (2008), Buron and Euzet (2005), Goertz (2004), Cheung (1993), Chisholm (1993), Olson et al. (2010) |
|  | *Dasyatis say* | P(1), O(1), | Amin (1998), Benz and Bullard (2004), Bullard et |
|  |  | A(5), M(1) | al. (2004), Campbell and Beveridge (1996), Cheung (1993), Olson et al. (2010) |
| Ginglymostomatidae | *Ginglymostoma cirratum* | A(9), O(6),<br>P(2) | Adamson and Caira (1991), Argumendo (1997), Benz and Bullard (2004), Borucinska and Caira (1993), Bruce and Cannon (1993), Bruce (1994), Caira and Gavarrino (1990), Caira et al. (1999), Caira et al. (2001), Caira et al. (2017), Cheung (1993), Credille et al. (1993), Garner (2013), Karns et al. (2017 unpublished), Olson et al. (2010), Palm (1995), Waldoch et al. (2010) |
| Gymnuridae | *Gymnura micrura* | A(1), P(2),<br>O(2), I(1) | Benz and Bullard (2004), Beveridge (1996), Campbell and Cheung (1993), Sarada et al. (1993) |
| Lamnidae | *Carcharodon carcharias* | P (5), I(1),<br>A(1), O(1) | Benz et al. (2003), Caira et al. (1999), Caira et al. (2017), Cheung (1993), Dippenaar et al. (2008), Hewitt (1979), Hogans and Dadswell (1985), Randhawa (2011) |
|  | *Isurus oxyrinchus* | A(6), P(1),<br>O(1) | Benz and Bullard (2004), Caira and Bardos (1996), Caira et al. (1999), Cheung (1993), Knoff et al (2002), Olson et al. (2010), Palm and Beveridge (2002), Ruhnke (1993), Williams (1978) |
| Myliobatidae | *Aetobatus narinari* | P(10), O(5),<br>I(1), A(1) | Al Kawari et al. (1996), Beveridge and Campbell (1988), Beveridge and Campbell (1998), Butler (1987), Caira et al. (1999), Caira et al. (2001), Caira et al. (2017), Campbell and Beveridge (2002), Cheung (1993), Ghoshroy and Caira (2001), Goertz (2004), Janse and Borgsteede (2003), Marie and Justine (2005), Morales-Serna et al. (2016), Olson et al. (2010), Whittington (1990) |
|  | *Manta birostris* | O(1), P(1) | Benz and Deets (1988), Cheung (1993) |
|  | *Rhinoptera bonasus* | A(7), O(3), | Benz and Bullard (2004), Borucinska and Bullard (2011), Caira et al. (2017), Cheung (1993), Garner (2013), Goertz (2004), Ivanov and Campbell |
|  |  |  | (2000), Olson et al. (2010), Stamper et al. (1998), Stidworthy (2017), Thoney and Burreson (1986) |
| Narcinidae | *Narcine bancroftii* | A(2) | Caira et al. (1999), Cheung (1993) |
| Pristidae | *Pristis pectinata* | A(1), O(2) | Benz and Bullard (2004), Cheung (1993), Kearns et al. (2010) |
| Rajidae | *Dipturus laevis* | A(1), O(1),<br>P(1) | Benz and Bullard (2004), Cheung (1993) |
|  | *Raja eglanteria* | A(1) | Becker and Overstreet (1979), Bruce (1994) |
|  | *Raja texana* | A(1) | Valdo-Zalik and Campbell (2011) |
| Rhincodontidae | *Rhincodon typus* | P(3) | Cheung (1993), Dyer (1988), Norman et al. (2000) |
| Rhinobatidae | *Rhinobatos lentiginosus* | A(2), O(1), | Anderson et al. (2012), Derome and Borucinska (2016), Valdo-Zalik and Campbell (2011) |
| Scyliorhinidae | *Scyliorhinus retifer* | A(2) | Cheung (1993), Khan and Newman (1981) |
| Sphyrnidae | *Sphyrna lewini* | O(7), A(5),<br>P(2) | Argumendo (1997), Benz and Bullard (2004), Beveridge and Campbell (1993), Bruce (1994), Caira et al. (2001), Healy (2003), Cheung (1993), Karns et al. (2017), Palm (1995), Palm (1999), Palm (2000), Palm and Beveridge (2002), Rokicki and Bychawska (1991), Sao Clemente and Gomes (1992), Shields (1985), Vassiliades (1985) |
|  | *Sphyrna mokarran* | A(2), O(8),<br>P(3) | Argumendo (1997), Beveridge and Campbell (1989), Beveridge and Campbell (1993), Caira et al. (1996), Caira et al. (2001), Cheung (1993), Maranick et al. (2011), Olson et al. (2010), Palm (1995), Palm (1999), Palm (2000), Schmidt and Beveridge (1990) |
|  | *Sphyrna tiburo* | A(3), O(5) | Benz and Bullard (2004), Beveridge and Campbell (2010), Bullard et al. (2001), Cheung (1993), Derome and Borucinska (2016), Garner (2013), |
|  |  |  | Olson et al. (2010), Stidworthy (2017) |
| Squatinae | *Squatina dumeril* | O(1) | Cheung (1993) |
| Triakidae | *Mustelus canis* | A(6), O(7),<br>P(1), M(2), | Caira et al. (1999), Caira et al. (2001), Caira et al. (2017), Cheung (1993), Cisló and Caira (1993), Leibovitz and Leibovitz (1985), Palm (1995), Palm (1999), Garner (2013), Goertz (2004), Knoff et al. (2001), Vardo-Zalik and Campbell (2011), Vassiliades (1985) |
| Torpedinidae | *Tetronarce nobiliana* | A(2), O(1), | Cheung (1993), Llewellyn (1960), Vassiliades (1985) |

**Fig. 1** depicts the pathogens within this study by percentages of the whole (totaling 448) and readily shows that cestodes are nearly half (47.5%) of all identified. Copepods and Monogeneans represent 20.6% and 13.4% respectively. Pathogens representing less than 0.9% are not depicted in this chart to retain readability.

**Fig. 1.**
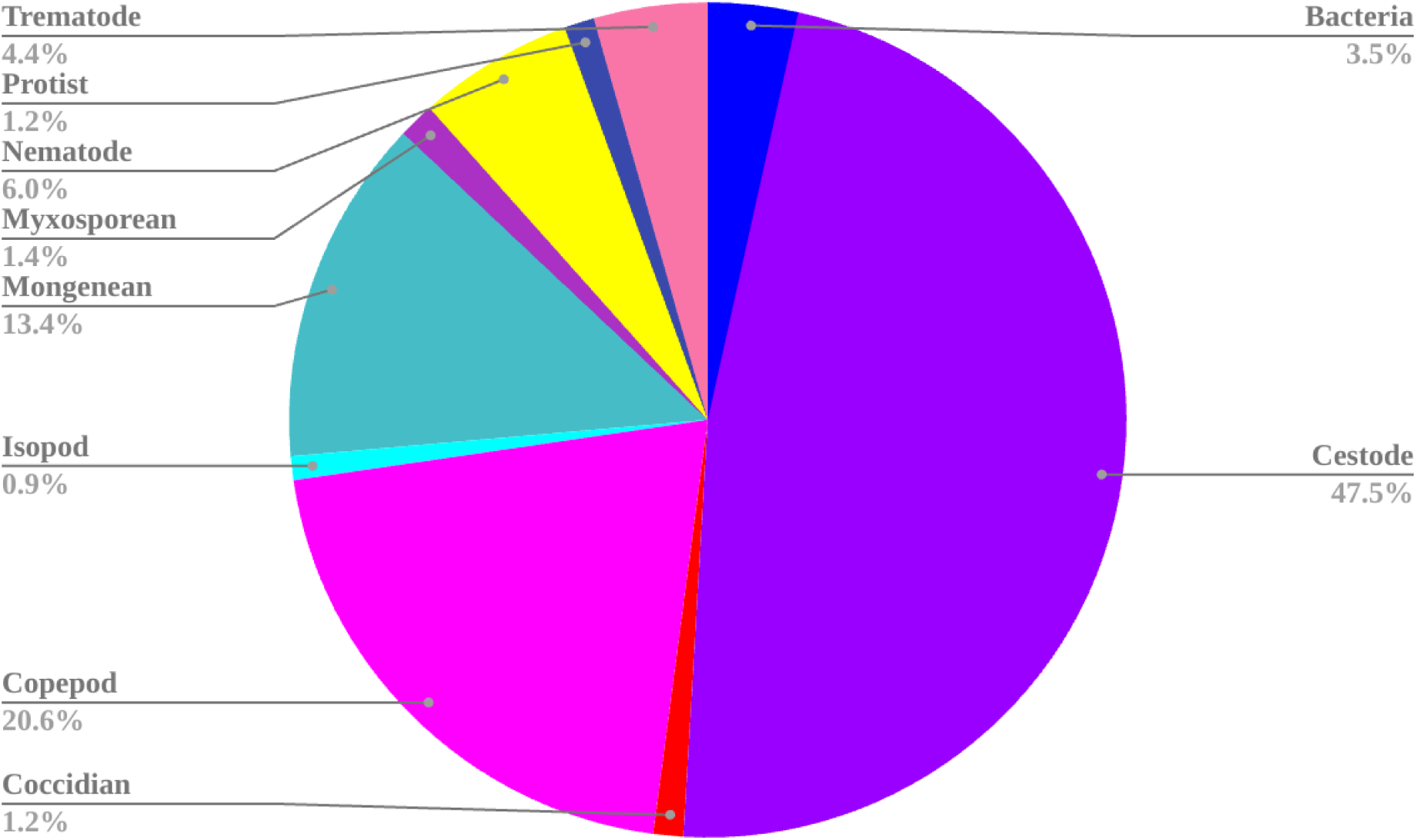
Reports of natural enemy type by percentage of the 448 identified pathogens.

Regression analysis found a significant positive correlation between the number of studies conducted on a given species and the number of natural enemies reported for that species (R squared = 0.492, p < 0.001, Df= 1,34; F=32.86). This relationship is depicted in **Fig. 2**. Regression also detected a significant relationship between the number of recorded prey items preferred by an elasmobranch species and the number of natural enemies (R squared= 0.13, p value = 0.026, df= 1 and 34; F=5.457). This relationship is depicted in the following **Fig. 3**.

**Fig. 2.**
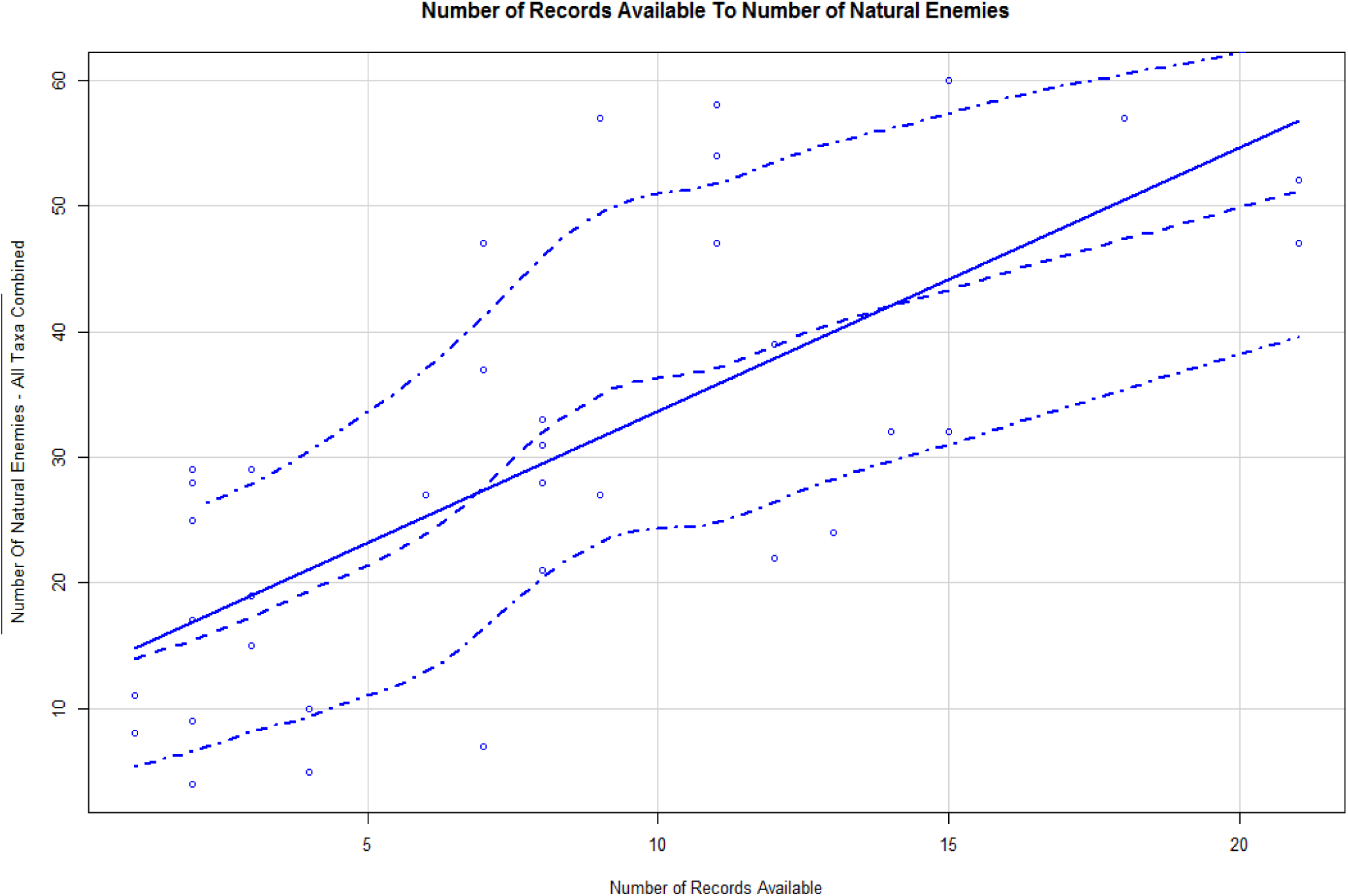
Linear Regression comparing the number of records available of each elasmobranch to the number of natural enemies. (R squared = 0.67, p-value <0.001, d.f. = 1,34; F = 67.64). There is a significant relationship between the number of records available about an elasmobranch and the number of natural enemies found. Dotted lines are LOWESS smoothing lines used to show possible relationships within the data

**Fig. 3.**
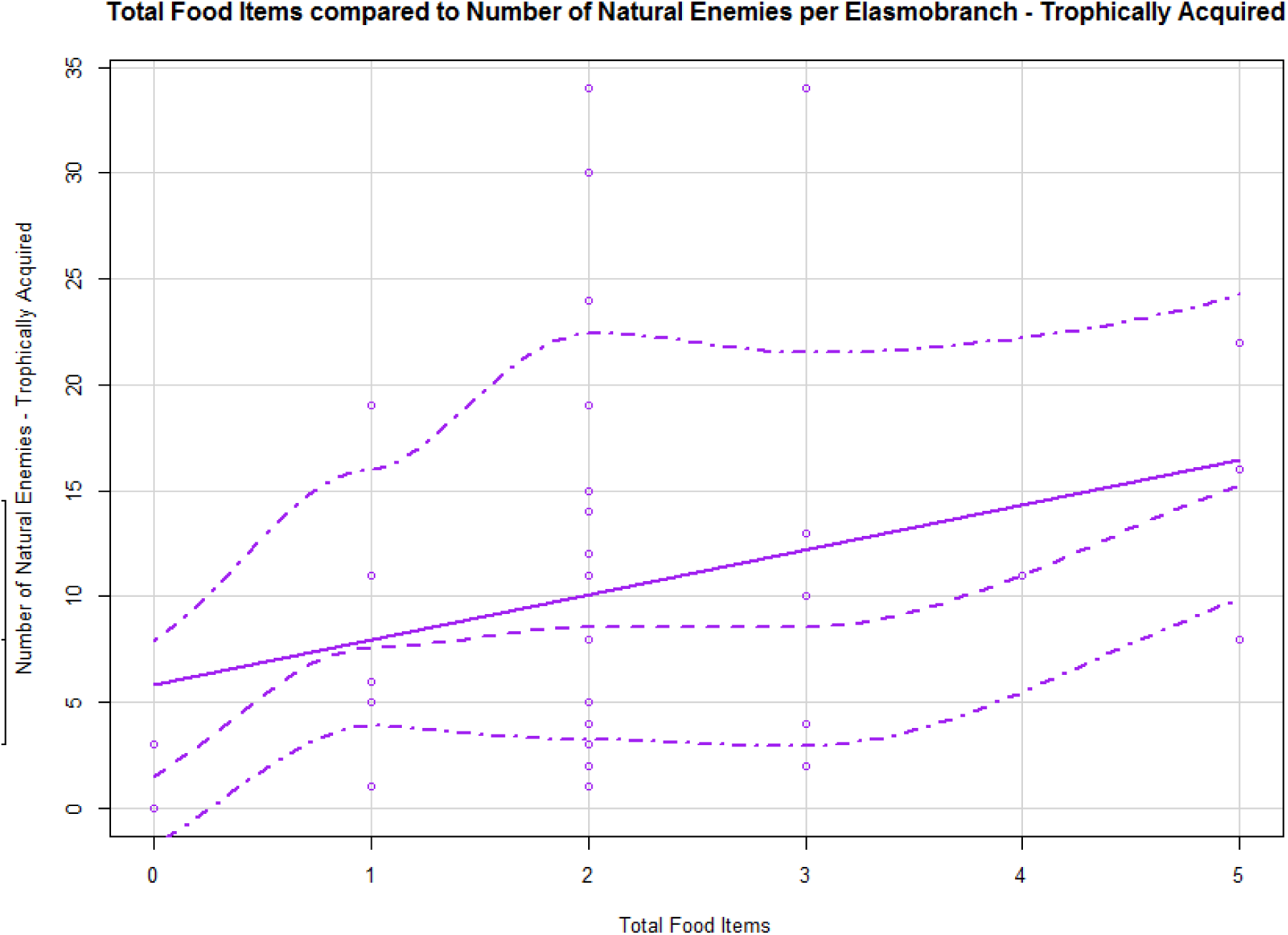
Linear regression between total food items and natural enemies trophically acquired. Dotted lines are LOWESS smoothing lines used to show possible relationships within the data

**Fig. 4** used the species level of elasmobranch with all natural enemies and clustered by using Bray-Curtis dissimilarity. A Bray-Curtis dissimilarity matrix ranges from zero to one, with zero indicating two samples as being identical. In **Fig. 4**, there seems to be some clustering due to *C. plumbeus* splitting the clusters. Similarly, **Fig. 5** depicts clustering by taxonomic Family using all natural enemies, where four clusters appeared.

**Fig. 4.**
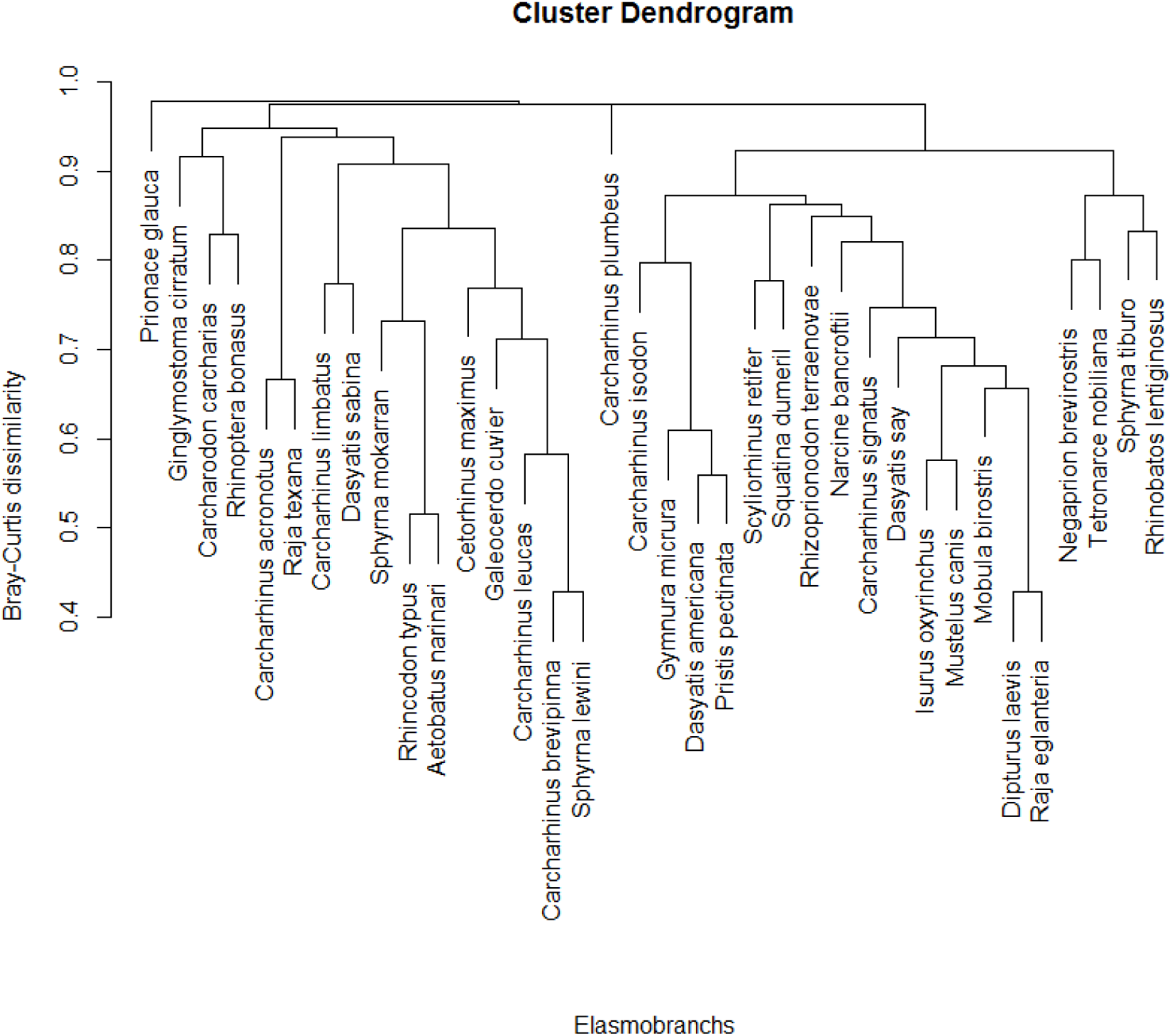
Species hierarchical cluster. No overt patterns appear, with the exception of the figure being split by *C. plumbeus*.

**Fig. 5.**
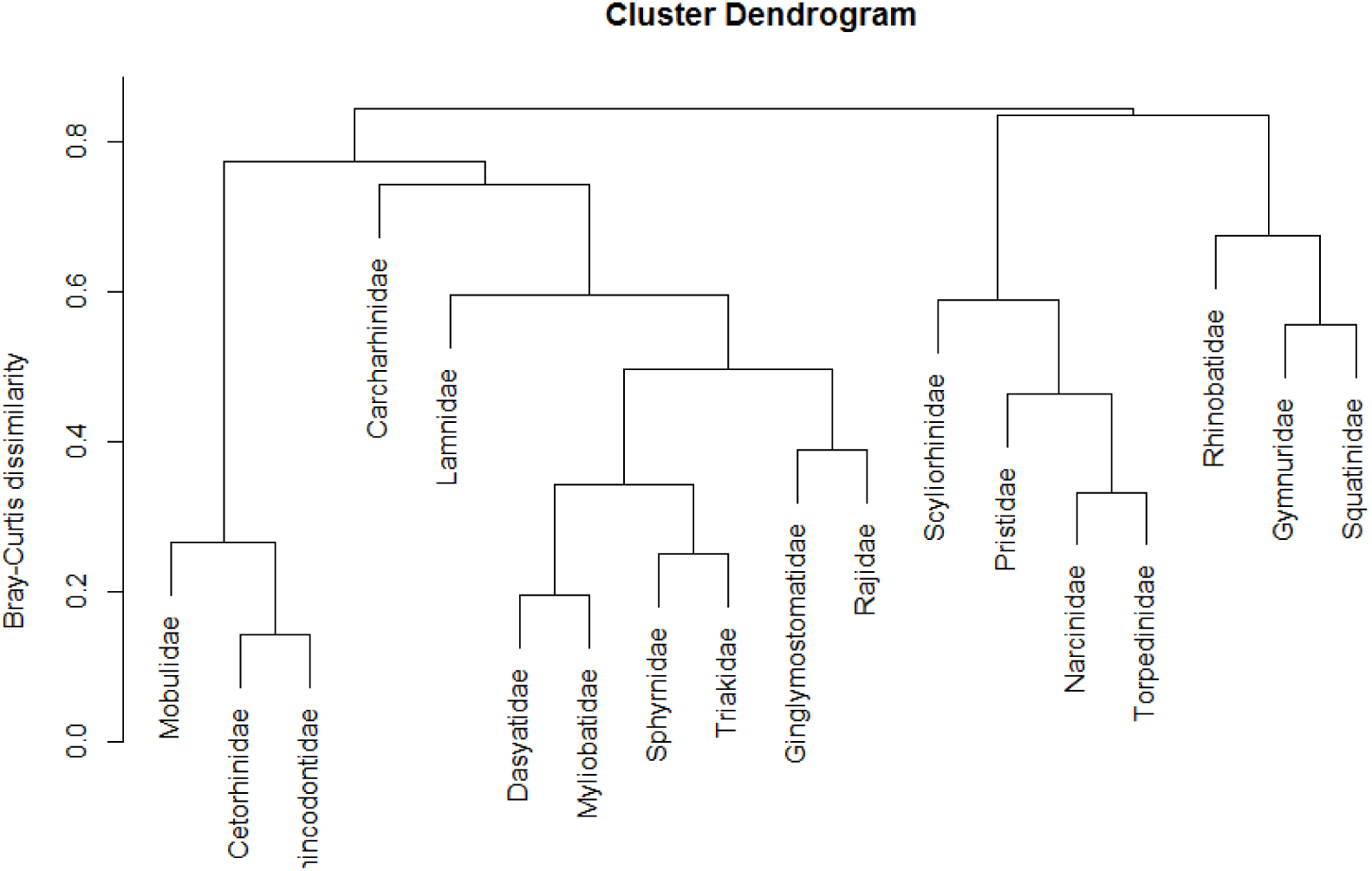
Hierarchical cluster by taxonomic family, split using natural enemy reports. There are about four clusters appearing.

The NMDS produced was used to examine the patterns between taxonomic order and natural enemy community similarity between samples. A cluster formed near the bottom of the figure as well as center-left between Triakidae, Scyliorhinidae, Rajidae, Dasyatidae, and Sphyrnidae. Rhinobatidae appeared alone near the top of the figure.

## Discussion

### Database Aggregation and Exploration

A statistically significant relationship was found between the number of natural enemies and number of records available. This indicates that with an increase in the number of publications related to natural enemies, there are new natural enemies described. The relationship indicates that the best way to learn more about elasmobranch disease and pathogens is to study them, with the follwing species being the most under studied: Carcharhinus isodon, Carcharhinus signatus, Rhincodon typus, Scyliorhinus retifer, Squatina dumeril, Gymnura micrura, Manta birostris, Narcine bancrofti, Pristis pectinata, Rhinobastis lentiginosus, Tetronarce nobiliana, Dipturus laevis, Raja eglanteria, and Raja texana.

In addition, there were differences in data available between the species reviewed (22 sharks, 11 rays, and three skates). For example, *Carcharhinus isodon* (Finetooth Shark), *Rhincodon typus* (Whale Shark), *Scyliorhinus retifer* (Chain catshark), and *Squatina dumeril* (Atlantic Angelshark) have less than four records documenting their pathogens. In comparison, the ray species of this study which are the most understudied include: *Manta birostris* (Manta), Narcine bancroftii (Lesser electric ray), *Rhinobatos lentiginosus* (Atlantic Guitarfish), and *Tetronarce nobiliana* (Atlantic torpedo). Each with two, one, three and three sources, respectively, documenting their pathogens and the diseases with which they are found. Lastly, all three species of skates included in the scope of this study have very few documented sources of their pathogens and diseases. The species are *Dipturus laevis* (Barndoor skate), *Raja eglanteria* (Clearnose skate), and *Raja texana* (Roundel skate). Each skate has less than three sources documenting natural enemies.

The highest number of reports per enemy type belong to the cestodes. Copepods, nematodes, trematodes, and monogeneans are the next most studied natural enemies.

### Bias in the Literature

Viruses, fungal infections, and bacteria are data deficient; there is little information on these pathogens infecting elasmobranchs. Availability and economic importance may play a role in the bias in host species studied (Baum and Worm 2009). Adamson and Caira (1991) noted that another reason for some elasmobranch species having less information may be due to the size of sharks and that performing thorough necropsies on sharks may not be a viable option. Anatomical areas that are not typically thought to harbor many parasites may be overlooked completely, such as the circulatory system. Al Kawari et al. (1996) explored helminths in the Arabian Gulf and noticed a bias toward studying larger organisms, species which do not require complex instrumentation, and those which are easiest to locate. This can lead to a vast underestimation of microorganisms in the less accessible species.

### Diet and Trophically Acquired Natural Enemies

There was a significant relationship between the number of trophically acquired natural enemies as well as number of prey items, however it is important to note that these analyses are being run with data that is fragmented due to species being understudied. Examination of stomach contents by necropsy of elasmobranchs may provide more information on trophically acquired parasites as well as the continued research into the spiral valve portion of shark digestive systems (Borucinska and Dunham 2000, Borucinska and Caira 1993, Cislo and Caira 1993, Curran and Caira 1995).

### Clustering of Elasmobranchs

**Fig. 4** did not display any tight clusters, this may be due to some elasmobranchs being more studied than others. In **Fig. 5**, the clusters are separated more clearly. These Families include the Basking shark, the Manta ray, and the Whale shark. These elasmobranchs are known to be more sluggish and slow moving individuals, which may explain why they would have similar natural enemies as each other. Carcharhinidae and Lamnidae clustered together. Both of these taxonomic Families include high energy, well studied shark species such as: Bull sharks, Blacktips, Sandbar sharks, White sharks, and the Shortfin Mako shark, etc.

This cluster has a number of well studied and under-studied species. Nurse sharks, Dusky Smoothhounds, Hammerheads and some rays have a large amount of information on natural enemies available. This cluster also includes skates. The second to last cluster includes Scyliorhinidae, Pristidae, Narcinidae, and Torpedinidae. The last cluster comprises: Rhinobatidae, Gymnuridae and Squatinidae. These final two clusters represent the least studied species, with the exception of the Rajidae family grouped elsewhere.

The generated NMDS is **Fig. 6** and had a very high r squared value (0.987, non-metric fit) indicating it was a good model to use for this data. The NMDS attempted to find patterns within the data, but the lack of information on Orders presents itself in the lack of very tight groupings. When more information on natural enemies is compiled and added to the database created in this study, a new NMDS should be run to look for new patterns based on dissimilarity between elasmobranchs.

**Fig. 6.**
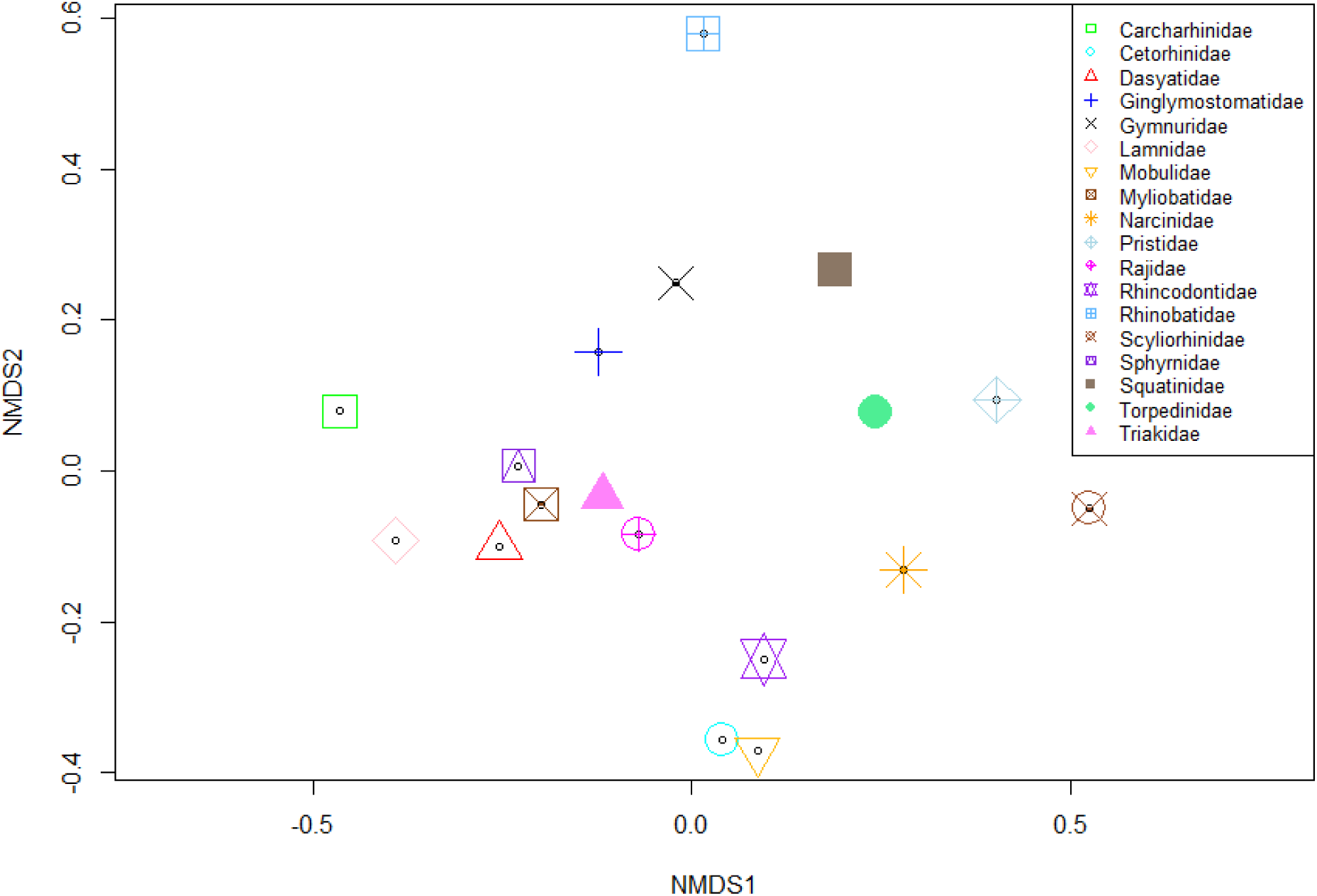
NMDS using elasmobranch taxonomic order. Some clustering appears, such as Cetorhinidae, Mobulidae, and Rhincodontidae near the bottom of the figure

The host or natural enemy list compiled in the present study will be of interest to resource managers and wildlife biologists who study elasmobranchs, providing much-needed information on potential natural causes of morbidity and mortality in local shark populations. This meta-analysis has provided further insight into the broad factors that shape natural enemy communities in these fish. Knowledge gaps focusing on individual elasmobranch species have been investigated and identified, which may open the door to further research.

## Summary and Conclusions

The above mentioned data deficient species would benefit from increased study by researchers and scientists in order to better understand and document the natural enemies of elasmobranch species, including those of commercial importance. With each new microbe description or recorded case of illness in an elasmobranch, the field of marine conservation gains insight into how to better conserve and protect at-risk species.

This literature review and meta analysis accomplished both research objectives of: creating the first comprehensive database of natural enemies from the selected Florida elasmobranchs, and the exploration of relationships between natural enemy community structure and life history traits, such as diet with a regression, hierarchical clustering, and a non-metric multidimensional scale model using taxonomy. Statistically significant relationships were found between the number of sources available and number of natural enemies present in an elasmobranch. The discovery of knowledge gaps pertaining to natural enemies and elasmobranch species has been uncovered, and discussion of underlying causes to bias leads to the indication that there is a lot of research needed in this field of study.

## Statements and Declarations

### Data Availability

The author confirms that all analyzed data for this article has been made available.

## Acknowledgements

This review was conceived after numerous marine microbiological experiences the author was fortunate enough to have and is grateful for the support from family and friends as well as mentors throughout this article’s development.

